# THE NEUROPROTECTIVE ROLE OF NICOTINE AND ASCORBIC ACID ON ALUMINIUM CHLORIDE (ALCL_3_)-INDUCED NEUROTOXICITY IN THE HIPPOCAMPUS OF ADULT MALE WISTER RATS

**DOI:** 10.1101/2023.08.09.552525

**Authors:** Kareemat Adebanke Akanbi, Aramide Ilesanmi

## Abstract

**Objectives:** Free radicals induced oxidative stress has been disrupted in the neurological functionality of the hippocampus in relation to aluminium chloride induced neurotoxicity. Ascorbic acid was been used as an antioxidants to prevent the oxidative damage induced by aluminium chloride. Therefore this present study was conducted to evaluate the antioxidant role Ascorbic acid and nicotine against the neurodegenerative consequences associated with aluminium chloride.

**Materials and method:** Thirty five (35) adult male Wister rats weighing 130-150g per body weight was used for this study. The rats were randomly grouped into five groups (A, B, C, D, and E) with each group having seven animals. Animals in group A(control group) were fed only water ad libitum for 21 days; animals in group B were administered Aluminium Chloride only with 100mg/kgBW for 21 days; animals in group C were administered aluminium chloride for the first 21 days with 100mg/kgBW, after which nicotine of 14mg/kgBW was administered. Animals in group D was administered with aluminium chloride of 100mg/kgBW for the first 21days after which Ascorbic acid was administered with 100mg/kgBW for the next 21days. Animals in group E were administered with Aluminium chloride, received 100mg/kgBW of Alcl_3_ for the first 21days then 14mg/kgBW Nicotine for 21days follow by administration of 100mg/kgBW of Ascorbic acid for the next 21days, concomitantly.

**Results:** Oxidative stress was observed from the degeneration the antioxidants (catalase, glutathione peroxidase, superoxide dismutase) in animals that were exposed to aluminium chloride. Histological examination showed that the hippocampus microarchitecture of animals exposed to aluminium chloride have a neurodegenerative changes characterized fragmentation, chromatolytic changes as well as some pyknotic changes in the purkinje and granule cell layers with a gross reduction in the cytoplasmic Nissl proteins. Administration of Ascorbic acid and nicotine was said to revert close to all the cytoarchitectural, biochemical, neurochemical, and immunohistochemical alterations induced by aluminium chloride neurotoxicity.

**Conclusion:** The results suggests that the administration of Ascorbic acid and nicotine may have caused a significant ameliorative effect against the neurotoxicity of aluminium chloride.

## INTRODUCTION

The human nervous system control behaviors, sensing ability and responding to external stimuli, it is responsible for mediating communication with the external environment; and it control the activities of all other organ systems. Thus it plays a cognitive role in maintaining metabolic balance and homeostasis (Everly and Lating, 2019).The consequences of damage to the nervous system can be recognize. Excessive damage to the nervous system can result in coma, convulsions, paralysis, dementia, strokes, and in coordination of the limbs. Even slight nervous system damage can impair reasoning ability, cause loss of memory, disturb communication, interfere with motor function, and impair health indirectly by reducing functions, such as attention and alertness, that ensure safety in the performance of daily activities(Everly and Lating,2019).

This research has been design to know the effects of nicotine and ascorbic acid on the neurotoxicity of the aluminium chloride on the brain. Ascorbic acid is an important metabolite for living things both at cellular, tissue and systemic levels. Ascorbic acid is a good reducing agent and facilitates many metabolic reactions and repair processes. Aluminium Chloride induced toxicity is implicated in the production of reactive oxygen and exhaustion of antioxidants leading to oxidative stress in brain. Nicotine and ascorbic treatment may therefore be capable of reversing the damage caused by aluminium chloride.

## MATERIALS AND METHODS

### Animal care and ethical approval

All protocols and treatment procedures were done according to the Institutional Animal Care and Use Committee (IACUC) guidelines and as approved by the Faculty of Basic Medical Sciences Ethics Review Committee, Osun State University, Nigeria. Thirty-five (35) adult male Wistar rats with an average weight of 130 to150g were purchased and housed in the animal house of the Faculty of Basic Medical Sciences, Osun State University. They were fed with rat pellet (purchased from Topfeed Feedmill, Osogbo) and water *ad libitum*. The animals were allowed to acclimatize for 1 week.

### Procurement of Materials and Preparation of Treatment Solutions

Crystalline salts of AlCl_3_ was obtained from Sigma Aurich and was dissolved in distilled water and adjusted to pH 7.4 with 0.1 M phosphate-buffer saline (PBS). Ascorbic acid (vitamin c) salt was also obtained from Sigma Aurich and was dissolved in distilled water. Pure Nicotine was procured from Sigma Aurich. and was siphoned using a syringe. These solutions were freshly prepared each morning of administration and kept at 4 ° C before use.

### Animal Grouping and Treatment

The thirty five (35) adult male rats were randomly assigned into 5 groups (A –E), each consisting of 7 rats (n = 7). The groups were treated as follows: Group A rats received distilled water for 21 days only. Group B animals were administered Aluminium Chloride only received 100mg/kgBW for 21 days. Group C were administered Aluminium chloride + Nicotene; received 100mg/kgBW of Alcl_3_ for the first 21 days then 14mg/kgBW was administered for the next 21days, respectively. Group D animals were administerd Aluminuim chloride + Ascorbic acid received 100mg/kgBW of Alcl_3_ for the first 21 days then 100mg/kgBW of ascorbic acid for the next 21 days, respectively. Group E animals were administered Aluminium chloride + Nicotene + Ascorbic acid received 100mg/kgBW of Alcl_3_ for the first 21days then 14mg/kgBW for 21days follow by administration of 100mg/kgBW of ascorbic acid for the next21days, concummitantly.

### Sacrifice and Sample Collection

On completion of treatments, rats for histological analysis were euthanized using 20 mg/kg of ketamine (intraperitoneal). Transcardial perfusion was done by exposing the left ventricle and injecting 50 ml 0.1 M PBS (pH 7.4) followed by 400 ml 4% paraformaldehyde (PFA). Excised brains were then rinsed in 0.25 M sucrose 3 times for 5 minutes each and then post fixed in 4% PFA for 24 hours before being stored in 30 % sucrose at 4 °C until further processing. Rats for enzymatic assays were sacrificed by separating the head from the trunk, to avoid the interference of ketamine with biochemical redox; brains were then excised, rinsed in 0.25 M sucrose 3 times for 5 minutes each and placed in 30 % sucrose in which they were stored at 4°C. Coronal sections of the hippocampus were obtained stereotaxically from each brain and these sections were later processed for different biochemical examinations.

### Histological techniques

Routine histological processing using Hematoxylin and Eosin staining method was carried out. Hippocampal sections were fixed in 4% buffered formosaline, dehydrated in ascending grades of alcohol, cleared in xylene, and infiltrated in molten paraffin wax before finally embedded in molten paraffin wax to form block. The paraffin block containing the tissue was then sectioned by the rotary microtome at 4 μm thickness. The sections were then floated in water bath at 40°C and transferred to a glass slide and stained with hematoxylin and eosin stains and cresyl fast violet for general neural architecture and nissl granulation following standard routine laboratory process. The slides were then viewed under light microscope and photomicrographs were taken at different magnifications.

### Biochemical Assays

Determination of Superoxide dismutase (SOD), Lipid peroxidation (MDA), Glutathione Peroxidase (GPx), Lactase Dehydrogenase (LDH), Nitric Oxide (NO), and Acetylcholine esterase (ACHE) activities was carried out on the homogenized hippocampi tissue of treated rats using spectrophotometric technique.

### Morphometric analysis

The body weights of the animals were taken on the first day of the experiment and the day of sacrifice. The weight gain was estimated as the difference between the initial weight of the animal and the final weight.

### Statistical analysis

All quantitative data were analyzed using GraphPad Prism® (version 6) software. Body weight, biochemical, neuroendocrine and immnunohistochemical outcomes were plotted in ANOVA followed with Tukey’s multiple comparisons test. Significance was set at p<0.05 (95% confidence interval). The results were represented in bar charts with error bars to show the mean and standard error of mean respectively.

## RESULTS

### Morphological Observations

Figure 1 shows the evaluation of the body weights of the animals taken on the first day of administration and the day of sacrifice. The weight gain was estimated as the difference between the initial weight of the animal and the final weight. Experimental animal in group B showed significant decrease in body weight gain in comparison to the control animals (group A) at p<0.001.Animals in group C showed significance decrease when compared to group A at the value of p<0.05 Animals post-treated with ascorbic acid and nicotine (groups C-E) had positive weight gain which were significant statistically when compared with those treated with AlCl_3_ only (Group B).

**Figure 1:**
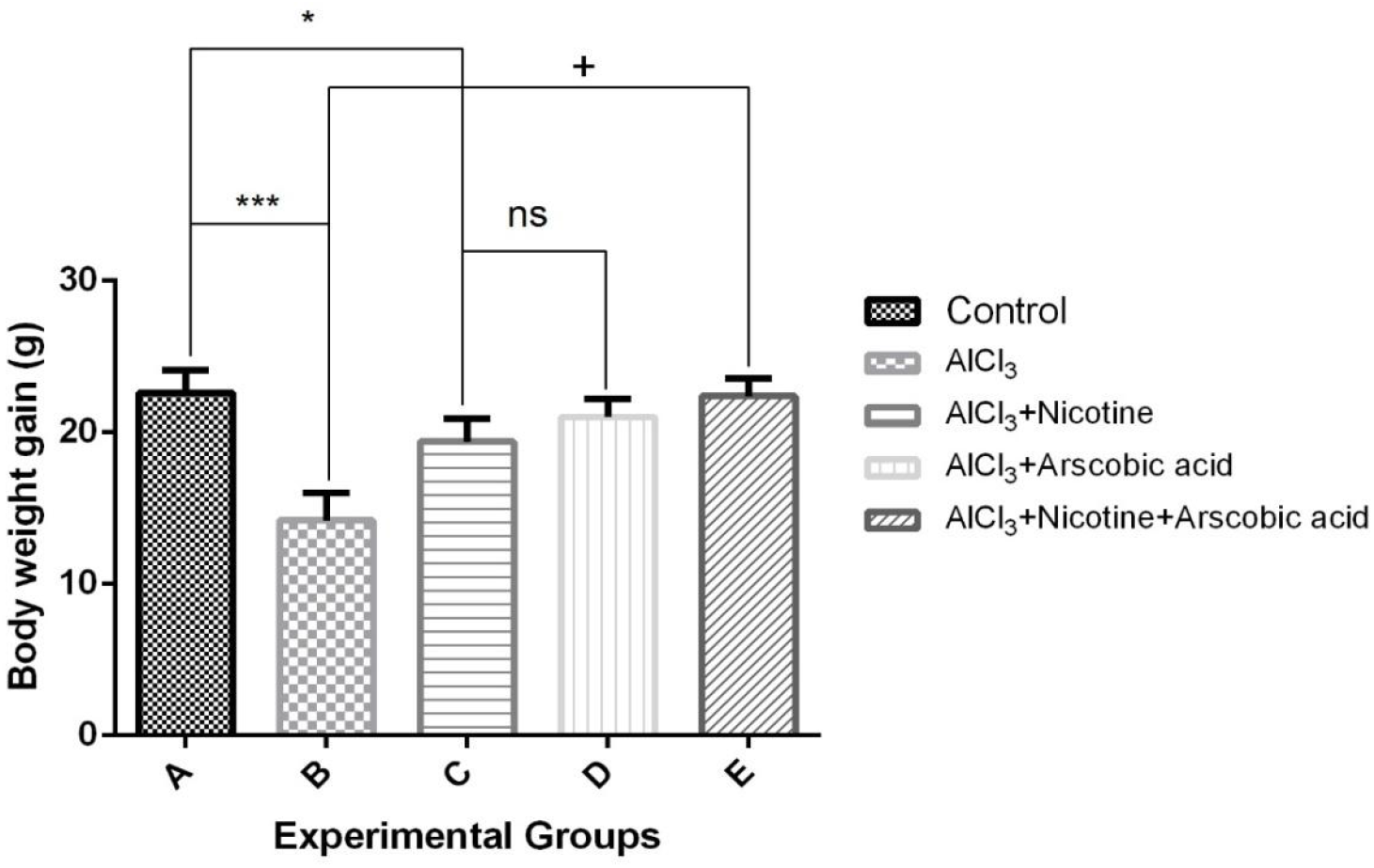
Bar chart showing weight of rats over the period of treatment, A= control, B= AlCl_3_, C= AlCl_3_+Nicotine, D= AlCl_3_+Ascorbic acid, E= AlCl_3_+Nicotine+Ascorbic acid. *is the significant level of difference in comparison to the control group (Group A) while **+** is the significant level of difference amongst other groups. **ns** means not significant.*/+ p< 0.05; **/++ p<0.01; ***/+++ p< 0.001.

### Biochemical Analysis

#### Glutathione Peroxidase

Figure 2 shows the analysis of the oxidative stress markers by GPx on the experimental animals. Group B (AlCl_3_) showed a significant decrease at p<0.001 of Gpx compared to the control group (groupA). Animals post-treated by nicotine (group C), showed a decrease at p<0.01 of GPx. Animals in group E showed increased significant level of GPx when compared to group B. Comparism of glutathione peroxidase levels between animals in group A and E, showed no significant difference in the level of GPx.

**Figure 2:**
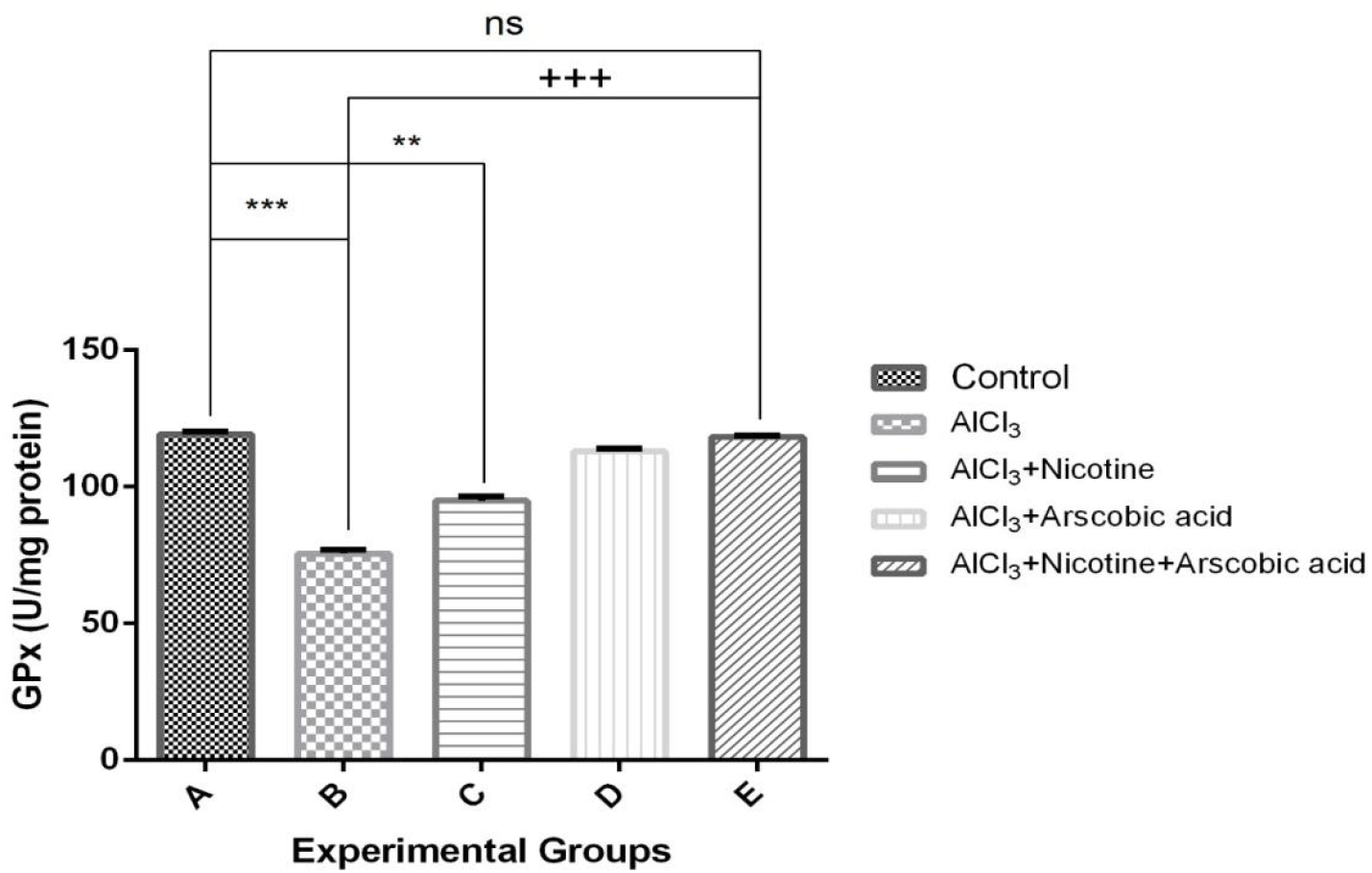
Bar chart of glutathione peroxidase in the hippocampus of experimental animals. A= control, B= AlCl_3_, C= AlCl_3_+Nicotine, D= AlCl_3_+Ascorbic acid, E= AlCl_3_+Nicotine+Ascorbic acid. *is the significant level of difference in comparison to the control group (Group A) while **+** is the significant level of difference amongst other groups. **ns** means not significant.*/+ p< 0.05; **/++p<0.01***/+++ p< 0.001.

### Superoxide Dismutase

Figure 3 showed the analysis of oxidative stress marker by SOD. Experimental animals in group B (AlCl_3_) showed high significance statistically at p<0.001 when compared to animals in group A (control). Animals in group A (control) showed increased significant statistically at p<0.001 when compared to group C (nicotine). Animals in group E showed high significance statistically at p<0.001 when compared to animals in group C. Animals in group B (AlCl_3_) and group C (nicotine) showed no significance.

**Figure 3:**
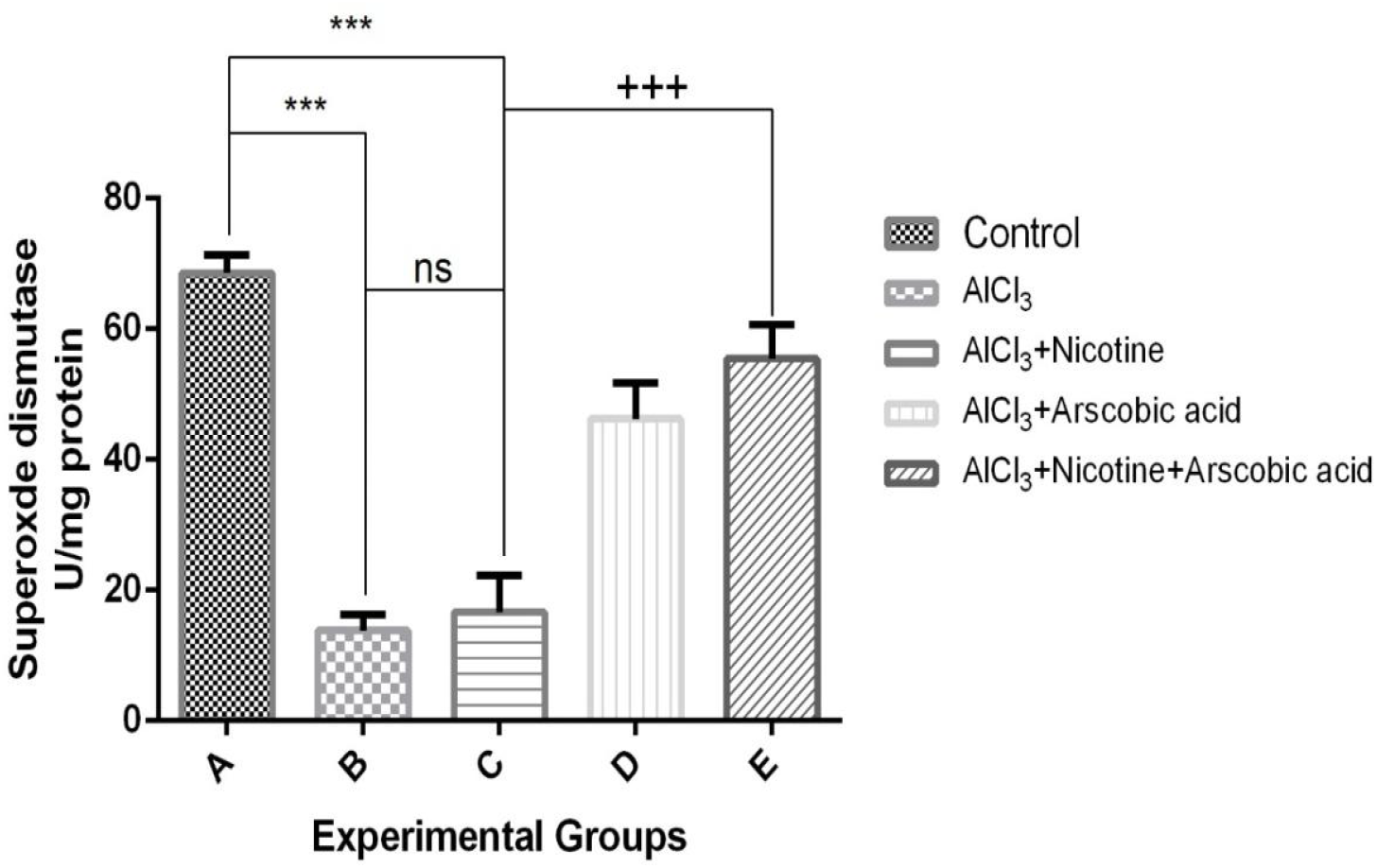
Activities of superoxide in the hippocampus of experimental animals. A= control, B= AlCl_3_, C= AlCl_3_+Nicotine, D= AlCl_3_+Ascorbic acid, E= AlCl_3_+Nicotine+Ascorbic acid. *is the significant level of difference in comparison to the control group (Group A) while **+** is the significant level of difference amongst other groups. **ns** means not significant.*/+ p< 0.05; **/++p<0.01;***/+++ p< 0.001.

### Lipid Peroxidation

The main effect of AlCl_3_ is to induce oxidative stress to the experimental animals. Figure 4 showed the level of neurotoxic effect of AlCl_3_ on the animals. Experimental animals in group B (AlCl_3_) showed high level of lipid peroxidation at p<0.001when compared to animals in control group (A). Animals in group C (nicotine) showed increased significance at p<0.001 when compared to group A (control). Animals in group C (nicotine) showed a decreased significance at p<0.01 when compared to group B (AlCl_3_)

**Figure 4:**
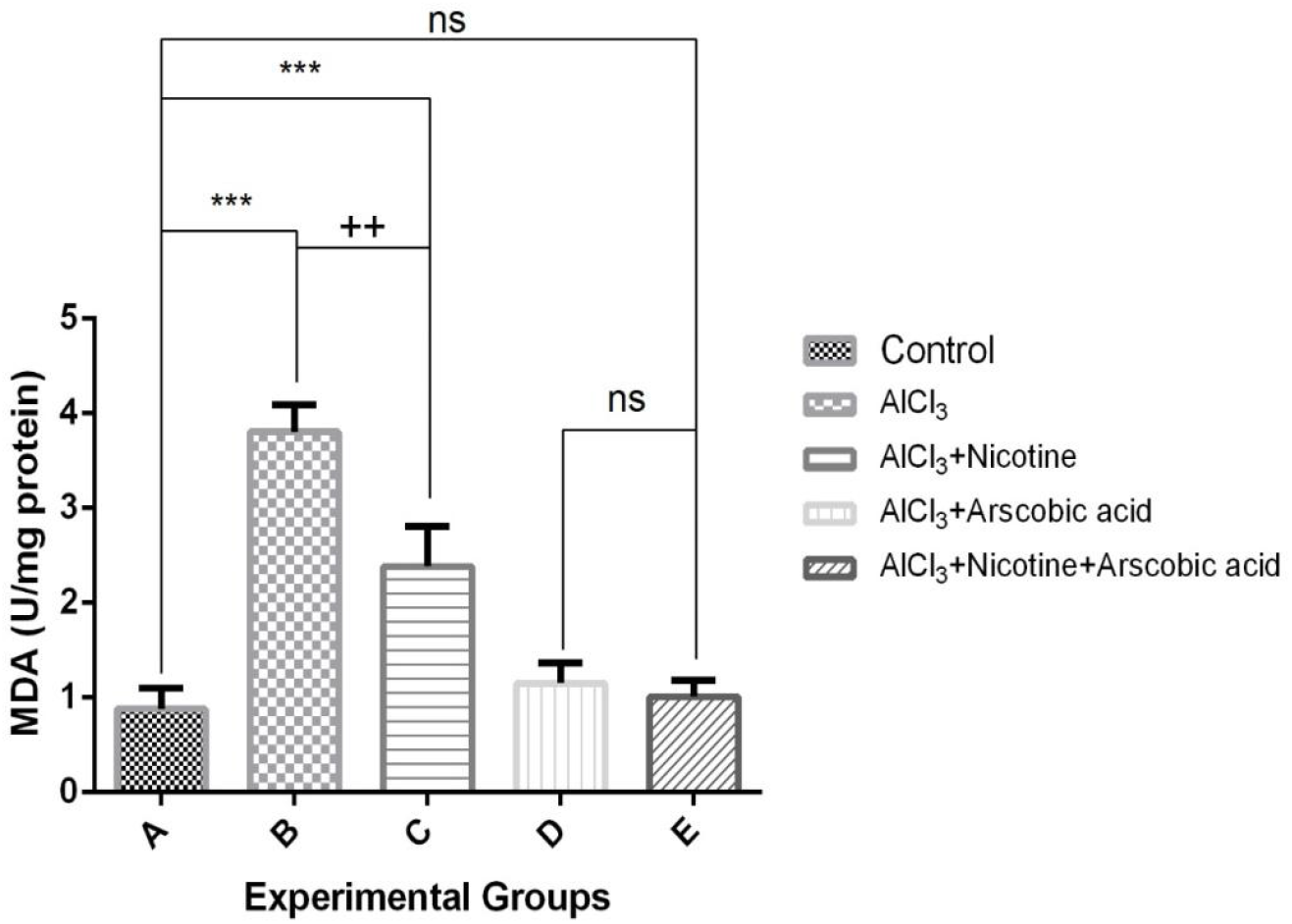
Lipid peroxidation levels in the hippocampus of experimental animals. A= control, B= AlCl_3_, C= AlCl_3_+Nicotine, D= AlCl_3_+Ascorbic acid, E= AlCl_3_+Nicotine+Ascorbic acid. *is the significant level of difference in comparison to the control group (Group A) while **+** is the significant level of difference amongst other groups. **ns** means not significant.*/+ p< 0.05; **/++ p<0.01; ***/+++ p< 0.001.

### Nitric Oxide

Figure 5 showed the enzymatic expression of Nitric Oxide on the experimental animals. Experimental animals in group B showed high significance statistically at p<0.001 when compared to group A (control). Animals in group C showed high significance statistically at p<0.001 when compared to group A. Animals in group E (nicotine +ascorbic acid) showed a high decreased significant at p<0.001 when compared to group B (AlCl_3_). Animals in group B and C showed no significance. Animals in group D and E showed no significance.

**Figure 5:**
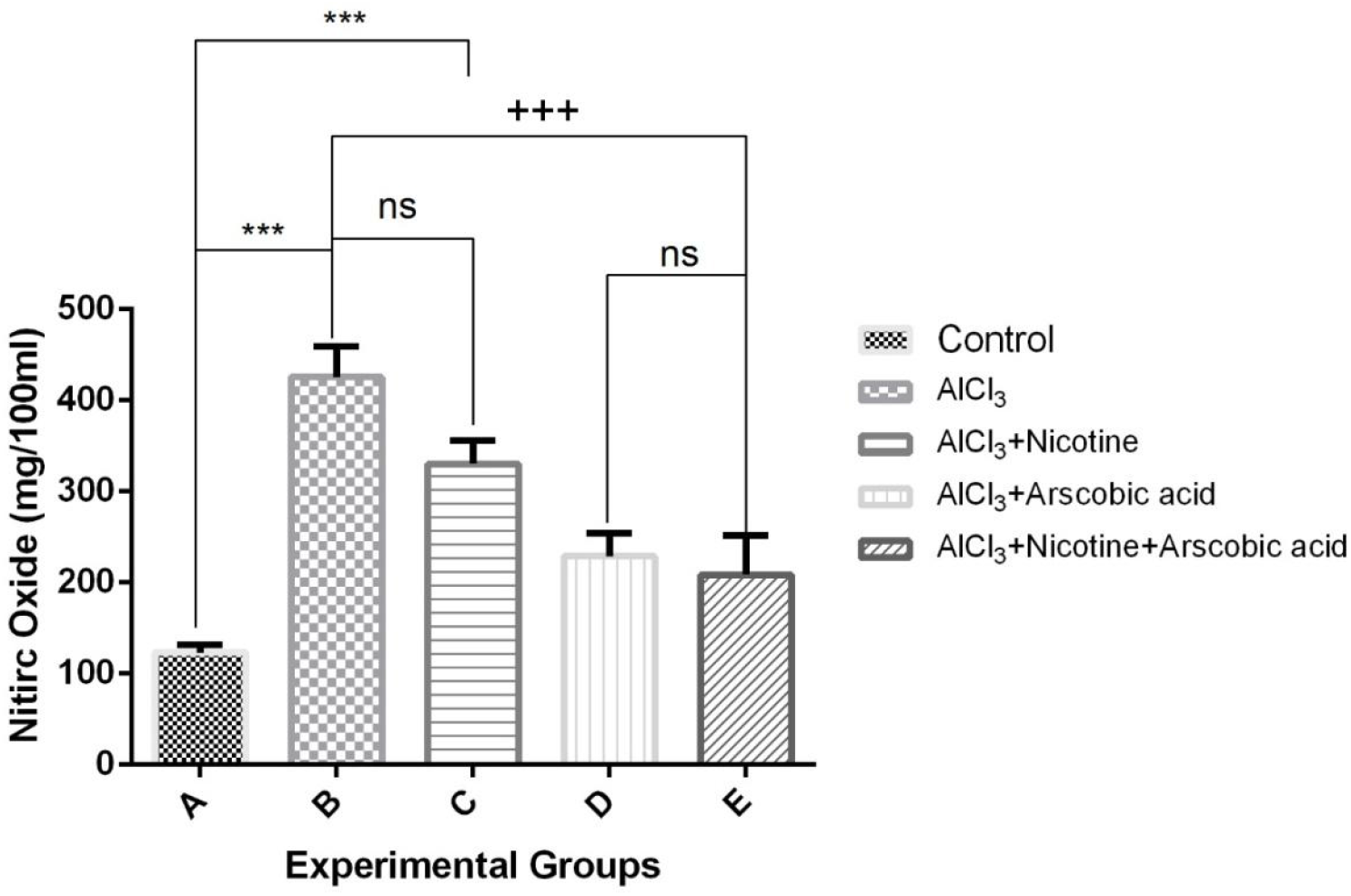
Activities Nitric Oxide in the hippocampus of experimental animals. A= control, B= AlCl_3_, C= AlCl_3_+Nicotine, D= AlCl_3_+Ascorbic acid, E= AlCl_3_+Nicotine+Ascorbic acid. *is the significant level of difference in comparison to the control group (Group A) while **+** is the significant level of difference amongst other groups. **Ns** means not significant.*/+ p< 0.05; **/++ p<0.01; ***/+++ p< 0.001.

### Acetylcholinesterase

The excitability of neurons in the cells of experimental animals were analyzed using the enzyme acetylcholinesterase. Figure 6 showed the activity of ACHE in the cells of hippocampus of the experimental animals. Animals in group A (control) and C (nicotine) showed no significance. Experimental animals in group B showed high level of significance at p<0.001 when compared to the control group (group A). Animals in group B showed high significance at p<0.01 when compared to animals in group C. Animals post-treated with ascorbic acid showed decrease level of AChE. Animals in group E (nicotine +ascorbic acid) showed no significance when compared to group C.

**Figure 6:**
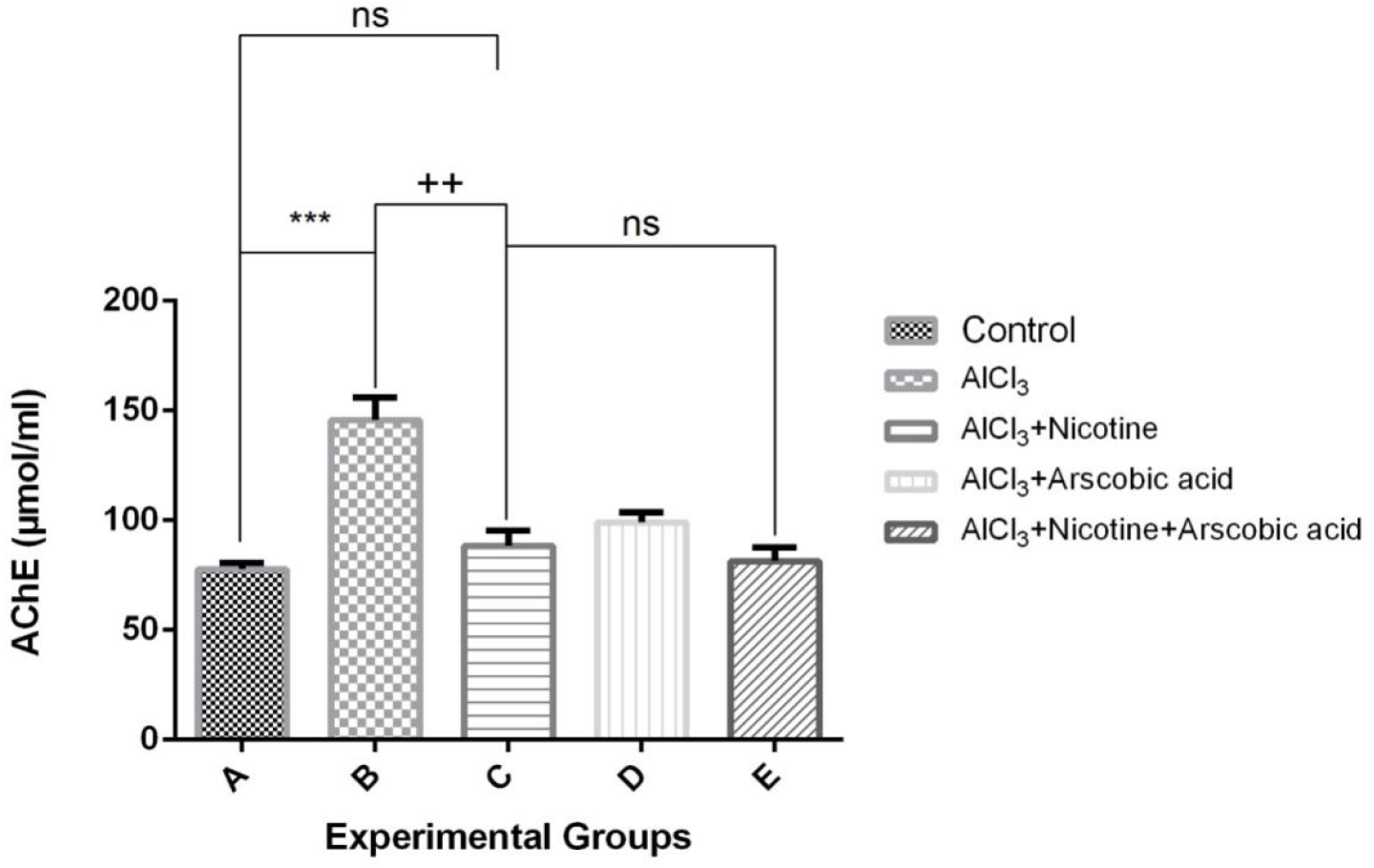
Bar chart showing activities acetylcholinesterase in the hippocampus of experimental animals. A= control, B= AlCl_3_, C= AlCl_3_+Nicotine, D= AlCl_3_+Ascorbic acid, E= AlCl_3_+Nicotine+Ascorbic acid. *is the significant level of difference in comparison to the control group (Group A) while **+** is the significant level of difference amongst other groups. **Ns** means not significant.*/+ p< 0.05; **/++ p<0.01; ***/+++ p< 0.001.

## Histological observations

### General Histology (Hematoxylin and Eosin Stain)

Representative micrographs of H and E staining showing the general cytoarchitecture of the hippocampus in Wistar rats. The control animals (group A) showed no observable altered panoramic morphological presentation of the hippocampal layers from this study across the various exposures and magnification (X100 and X400). In this group, the well outlined array of cells within the hippocampus can be seen distinctly arranged from the cornuammonus (CA) to the dentate gyrus (DG), (Figure 7). In addition, cellular density within this group appear normal across all hippocampal layers with appreciable spines and neuronal projections (black thin arrows), (Figure 7). AlCl_3_ treatment on the other hand, induced degenerative changes in the hippocampus characterised by fragmented pyramidal and granule cell layer with observable pyknotic cells (red thin arrows), (Figure 8). Also, there appeared to be a comparatively increased cell density in the hippocampal granular layer of this treated group.

**Figure 7:**
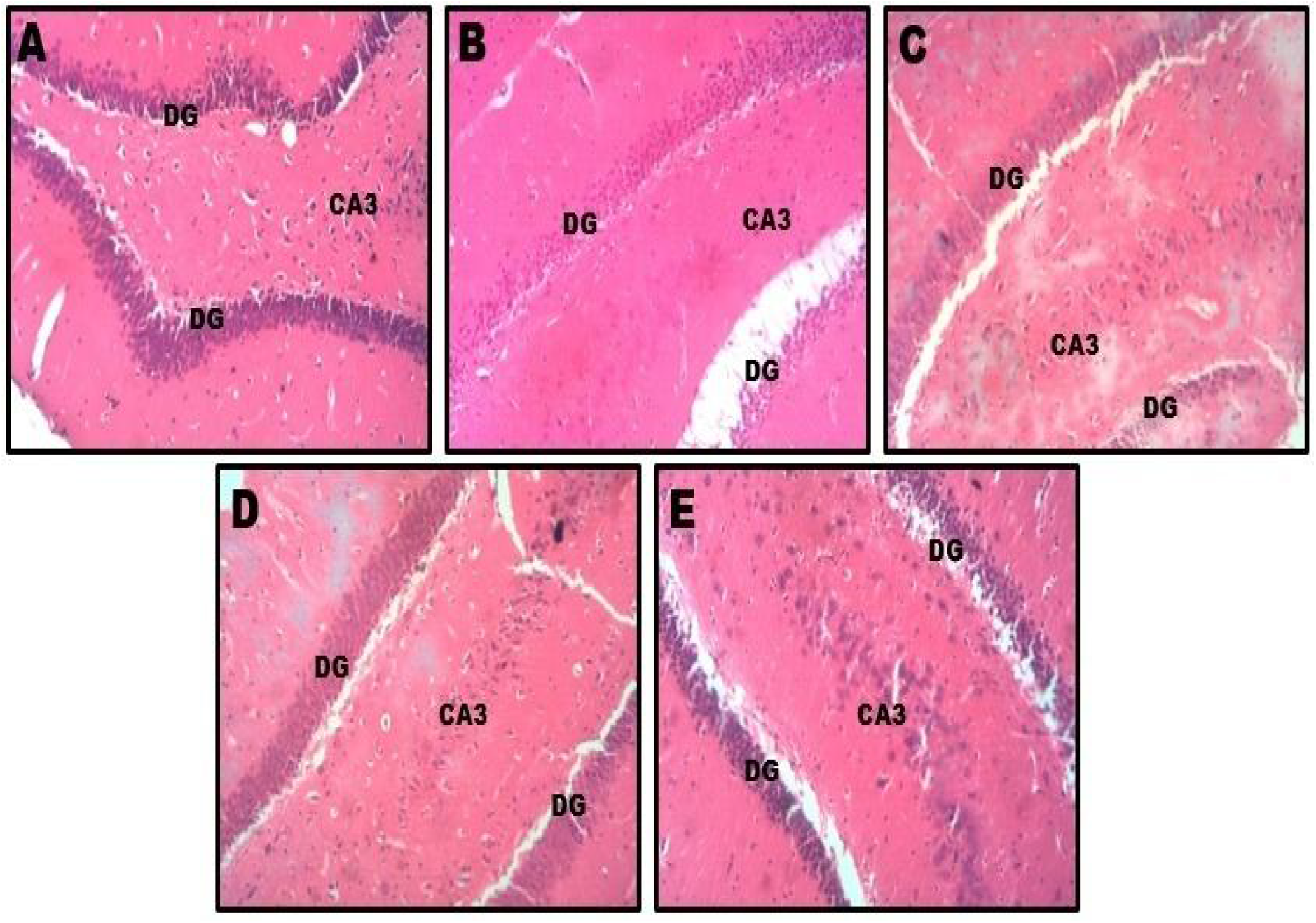
representative photomicrograph of the hippocampus showing the general histology at a panoramic view (H and E, x100). The Dentate gyrus (DG) composed of granule cells, Cornuamonus (CA3) containing pyramidal cells, are well demonstrated across the study groups.A= control, B= AlCl_3_, C= AlCl_3_+Nicotine, D= AlCl_3_+Ascorbic acid, E= AlCl_3_+Nicotine+Ascorbic acid.

**Figure 8:**
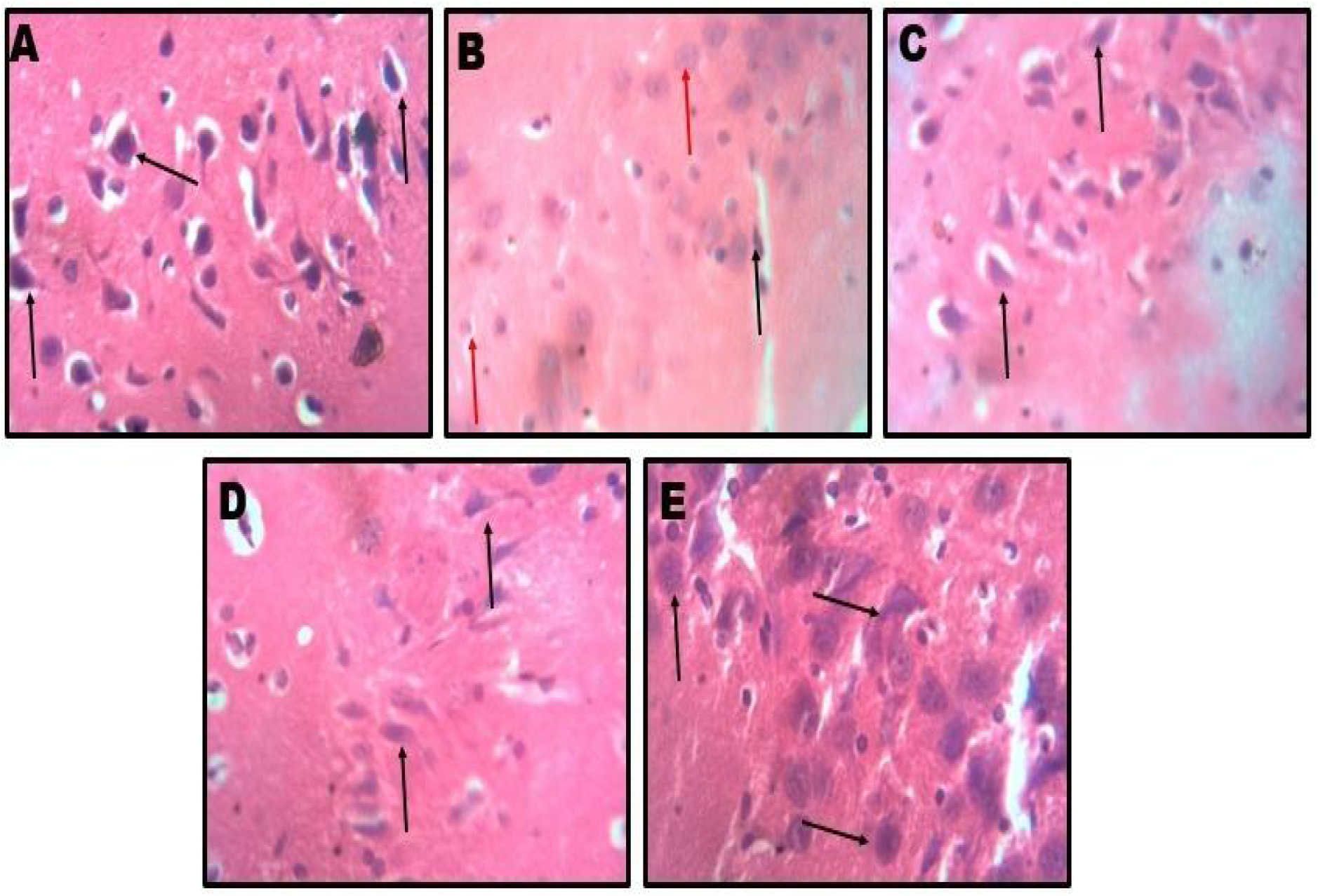
representative photomicrograph of the hippocampus showing the general histology higher magnification (H and E, x400) A= control, B= AlCl_3_, C= AlCl_3_+Nicotine, D= AlCl_3_+Ascorbic acid, E= AlCl_3_+Nicotine+Ascorbic acid.

Animals post-treated with nicotine (group C) showed slight degenerative changes (red thin arrows), (Figure 8) with overall similarity to the morphologic appearance of control group; post-treatment with ascorbic acid (group D) preserved the integrity of the hippocampus. The cortical layers of animals in this group seem better structured and delineated with distinct layering when compared to AlCl_3_ treated animals (group B). However, animals in group E that received combined administration of nicotine and ascorbic acid after treatment with AlCl_3_ had similar hippocampal microarchitecture to the control group. Photomicrographs showed no observable altered morphological presentation of the hippocampal layers.

### Nissl Profile

Histochemical demonstration of nissl profile by Crysl fast violet stain (CFV stain X400) (Figure 9) across hippocampal sections within the study groups showed normal expression of nissl protein in the control group (A) and in animals co-treated with nicotine and ascorbic acid (group E). Post-treatment with ascorbic acid only (group D) also showed a nissl profile consistent with the control group and characterized with normal and densely populated Nissl proteins, well stained and outlined neurons. AlCl_3_ (group B) caused severe chromatolytic changes as well as some pyknotic changes in the pyramidal and granule cell layers with a gross reduction in the cytoplasmic Nissl proteins (red arrow). Group C animals that were post-treated with nicotine only after treatment with AlCl_3_, showed reduced expression of nissl protein when compared with the control group (A) and co-treatment group (E).

**Figure 9:**
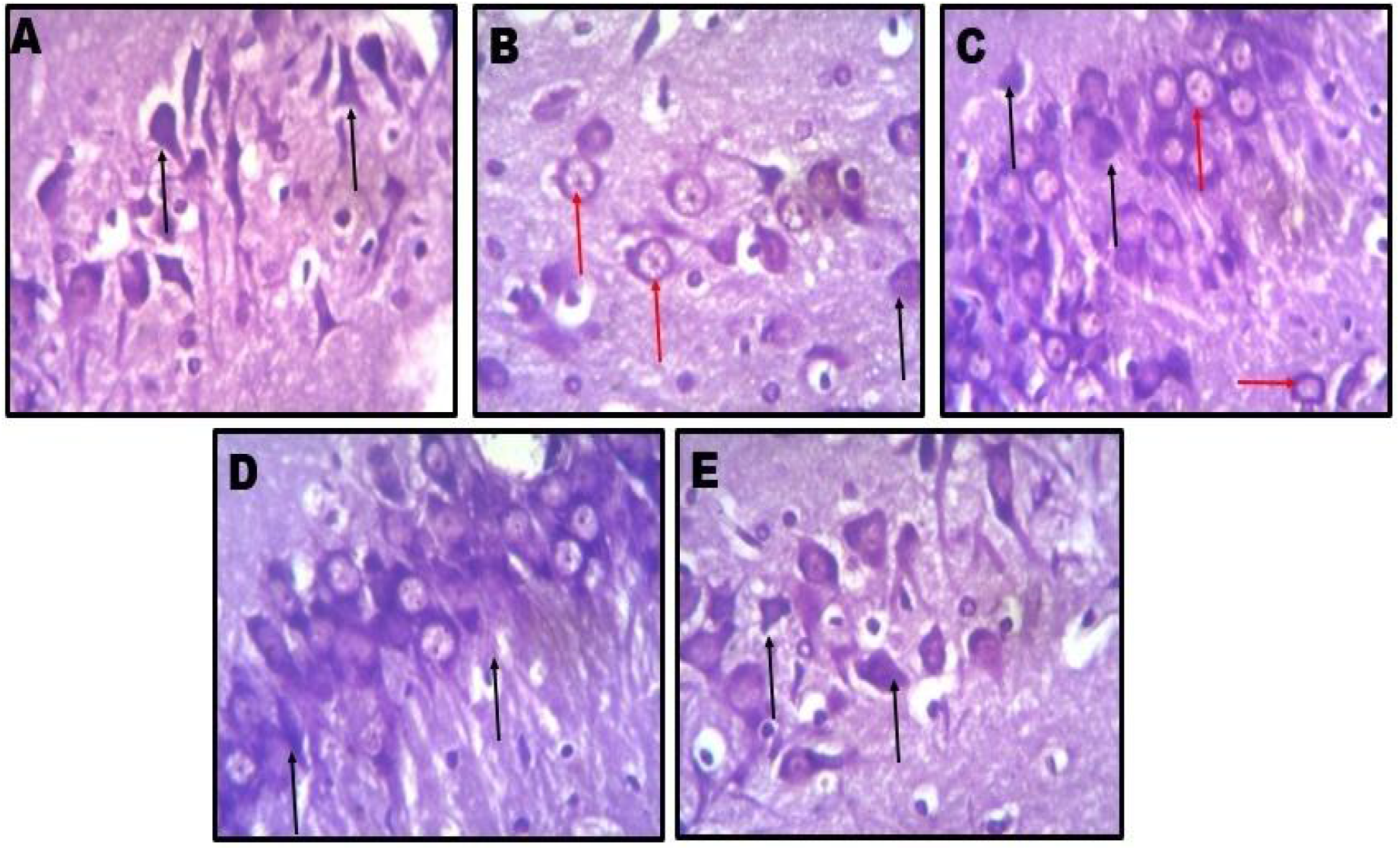
representative photomicrograph of hippocampus showing the Nissl substances at a high power magnification (crysl fast violet (x400).A= control, B= AlCl_3_, C= AlCl_3_+Nicotine, D= AlCl_3_+Ascorbic acid, E= AlCl_3_+Nicotine+Ascorbic acid.

### Immunohistochemistry

Evaluation of immunohistological expression of neurofibrilliary tangles using the congo red stain. In this study, there was no observable expression of neurofibrilliary tangles across the treatment groups.

## DISCUSSION

The following are the effect of AlCl_3_ on experimental animals with post-treatment of nicotine and ascorbic acid on the hippocampus of adult wistar rats.

The weight gain and loss were estimated as the differences between the initial weight of the animal and the final weight. Experimental animals in AlCl_3_ group (group B), showed a significant decrease in body weight gain in comparison to the control group animals (group A), this was in line with a previous study done by (Buraimoh and Ojo, 2014), which showed that aluminium salt have effect on drinking pathway in the brain of the rats, then results in decrease mitochondrial energy metabolism. Animals post-treated with ascorbic acid and nicotine (groups C-E) had a positive increase in weight gain which were significant statistically when compared with those treated treated with AlCl_3_ only (group B). Figure 1 shows the significant differences.

The histological structure of the hippocampus of the animals that were exposed to Aluminium chloride in this present study showed neurodegenerative changes and was characterized by fragmented pyramidal and granule cell layer with observable pyknotic cells. Comparative increased cell density in the hippocampal granular layers was observed in the treated group (group B). Animals post-treated with nicotine (group C), showed slight degenerative changes compare to the control group (group A), post-treatment with ascorbic acid (group D) preserved the integrity of the hippocampus. Animals in group E that receive the combined administration of ascorbic acid and nicotine after treatment with aluminium chloride had similar hippocampal microarchitecture compared to the control group.

Nissl profile demonstration by Cresyl fast violet stain (Fig 10) across hippocampal sections within the study groups showed normal expression of nissl protein in the control group (A) and in animals co-treated with nicotine and ascorbic acid (group E). Post-treatment with ascorbic acid only (group D) also showed a nissl profile consistent with the control group and characterized with normal and densely populated Nissl proteins, well stained and outlined neurons. AlCl_3_ (group B) caused severe chromatolytic changes as well as some pyknotic changes in the pyramidal and granule cell layers with a gross reduction in the cytoplasmic Nissl proteins (red arrow). Group C animals that were post-treated with nicotine only after treatment with AlCl_3_, showed reducedexpression of nissl protein when compared with the control group (A) and co-treatment group (E). Aluminium chloride has reported to be a widely spread neurotoxicant and industrial pollutant, which eventually leads to vital organ damages such as the liver, kidney, testes and brain (Squire *et al*.,2019).

**Figure 10:**
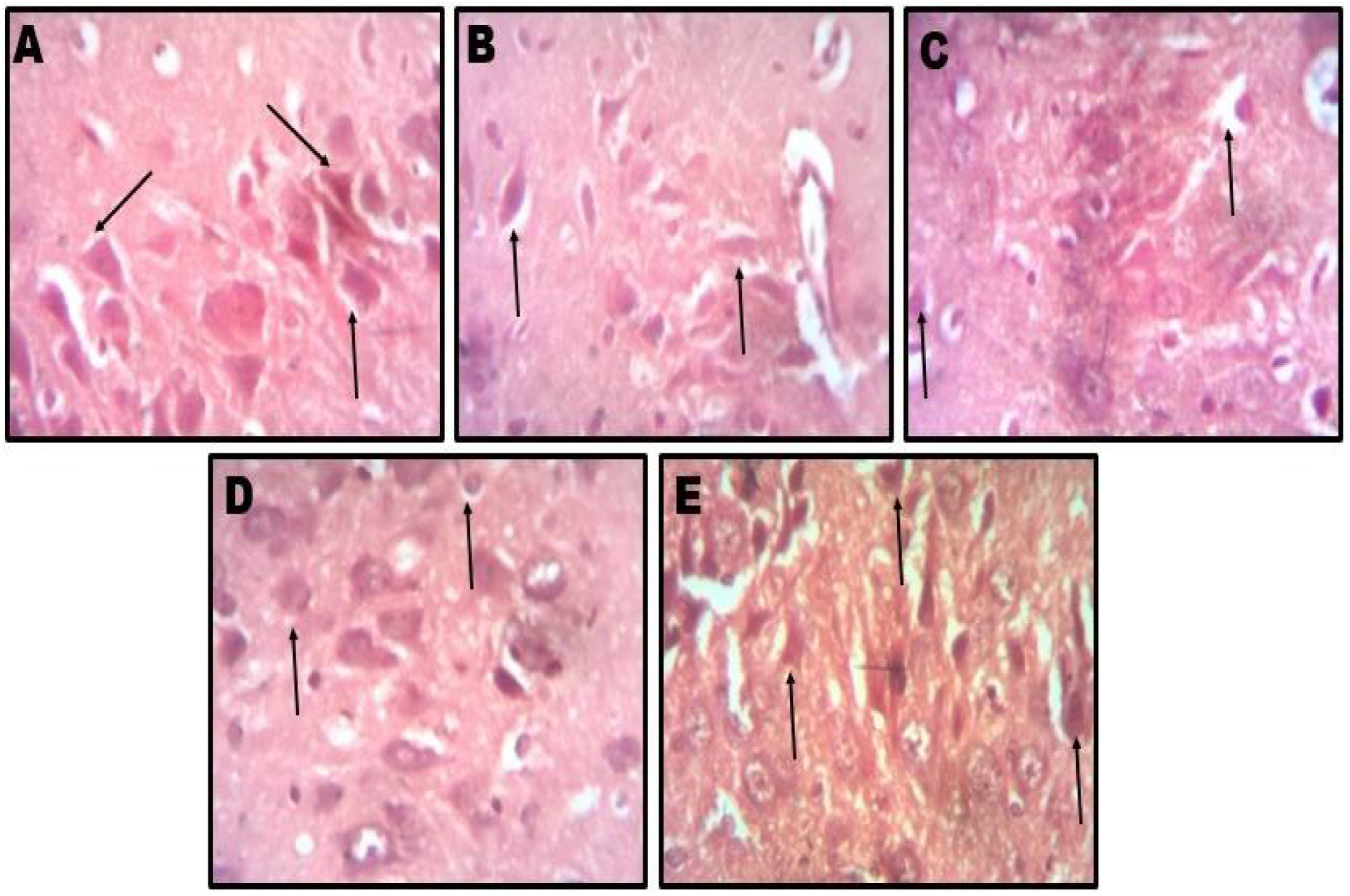
representative photomicrograph of hippocampus showing neurofibriliary tangles at a high power magnification (Congo red x400).A= control, B= AlCl_3_, C= AlCl_3_+Nicotine, D= AlCl_3_+Ascorbic acid, E= AlCl_3_+Nicotine+Ascorbic acid.

The major harmful effects of these heavy metals on human health are mediated via oxidative stress. The results of this present study showed the administration with aluminium chloride induces an increased level of lipid peroxidation product, MDA, in the blood of rats which were accompanied by increased formation of ROS, and inhibition of superoxide dismutase enzyme. The oxidative stress caused by the administration of aluminium chloride leads to the neuronal destruction in the hippocampus. Reactive oxygen species (ROS) are thought to be the major ones responsible for the alteration of macromolecules which is often termed oxidative stress (Hybertson *et al*., 2011). The lipid peroxidation and breakage of lipids with the formation of reactive compounds leads to changes in the permeability and fluidity of the membrane lipid bilayer and can dramatically alter cell integrity (Dix and Aikens, 1993).

This observation is in agreement with the previous investigation which showed chronic treatment with aluminium chloride caused dramatic encephalopathic morphopathological lesions (Abd-Elghaffar SKh, *et al*., 2005). Although, AlCl_3_ and nicotine showed a slight significant in the lipid peroxidation but this present study showed that ascorbic and nicotine (group D and E) helps to ameliorate the effect of the MDA in the hippocampus and helps to elevate the drastically decreased SOD, GSH, in the hippocampus of wistar rats exposed to aluminium chloride. This is in line with the previous investigation which showed that ascorbic acid detoxifies reactive oxygen species or repairs the resulting damages caused by free radicals (Hacisevki, 2009).

This present study also showed the significant role of superoxide dismutase, which showed increased in level of content in the control group (Group A) compared to the experimental group (group B). The difference between the AlCl_3_ group only (Group B) and AlCl_3_ + nicotine (Group C) is not significant, because both groups were dependent on the AlCl_3_ toxicity. Post-treatment with ascorbic acid and nicotine showed elevated effect to both group D and E. The ameliorative effect of SOD is to play a very important antioxidant defence against oxidative stress in the hippocampus (Younus, 2018).

This present study showed an increase in LDH content in the hippocampus of the experimental animals in group B and C. It was discovered that LDH caused tissue damage to the cells of hippocampus which indicate loss in membrane integrity. This was in agreement to a previous study done by (Fernanda *et al*., 2007), which suggested that high level of LDH is a marker of tissue necrosis, and a decrease of cell viability. Animals post-treated with ascorbic acid and nicotine (group D and E), showed a significance difference in LDH content, this shows that ascorbic acid acts as a anti-oxidant, contributing to the formation of hydroxyl radicals that may lead to lipid, DNA, or protein oxidation (Garcia, *et al*., 2017).

It was discovered in this study that AlCl_3_ treated group showed high significant in G-6-PDH content when compared to the control group (A), this resulted in mitochondrial dysfunction in the cells of the treated animals, this reduced the energy utilization in the treated animals. This in line with a previous study done by (Yusuf *et al*., 2017) which showed that aluminium treated group caused a significant degree of inhibition in the activity of G-6-PDH enzyme when compared to group A. Animals treated with AlCl_3_ and nicotine showed a moderate significance difference when compared to A. Ascorbic acid group showed a slight significant increase when compared to group C (nicotine). Ascorbic acid was discovered to be able to reversed the energy utilization in animals in group D by ameliorating the oxidative stress property of aluminium chloride in this study.

Nitric oxide was discovered to caused damages to the cells of the treated animals by inducing oxidative stress by aluminium chlorides. AlCl_3_ treated group showed a high significant level of NO when compared to the control group, nicotine group showed a moderate significance when compared to control group. Animals in group D and E showed no significance difference. This present study was in line with a previous done by Dawson *et al*., 1996, which suggested peroxynitrite and oxygen free radicals can be generated in excess of a cell antioxidant property resulting in severe damage to cellular constituents including proteins, DNA and lipids.

Reduced glutathione peroxidase (GPx) is a cytosolic enxyme that catalyzes the reduction of hydrogen peroxide to water and oxygen as well as catalyzing the reduction of peroxide radicals to alcohols and oxygen, which helps to prevent lipid peroxidation and maintain intracellular homeostasis as well as redox balance. Aluminium chloride treated group showed a significant decreased in GPx level when compared to control group. This is suggestive of generation of ROS as a result of mitochondrial dysfunction caused by AlCl_3_ administration. Weber *et al*., 1991, suggested that GPx deficiency contributes to seizure, as well as neurodegenerative disorders. Animals in group A and E showed no significant difference. The antioxidant property of ascorbic acid were able to ameliorate the effect of GPx in the hippocampus of animals in group D.

It was discovered in this present study that aluminium chloride treated group (B) showed increased significant in the content of acetylcholinesterase (AChE), this showed that there is over expression of AChE. Over expression of acetylcholinesterase in the cells of the treated animals results in dysfunction in neuronal function which leads to over excitation to the animals. This was in line with a previous study done by Mirjana *et al*., 2013, which suggested that AChE is involved in the termination of impulse transmission by rapid hydrolysis of the neurotransmitter acetylcholine. Animals in group C and E showed no significance difference.

## 5.2 CONCLUSION

In conclusion, the weight of the experimental animals, biochemical analysis, immunohistochemical examination, histological investigation results showed that there is neurotoxic effect of AlCl_3_ in the hippocampus. These results showed that induction with Aluuminium chloride had potentially deleterious effects on the brain thereby increasing oxidative stress level, over expression of enzymes such as LDH, MDA, alteration in the general cytoarchitecture of the hippocampus. However, post-treatment with ascorbic acid helped to ameliorate the effect of the degenerative impact of aluminium chloride in the hippocampus. Whereas, nicotine was observed to cause more deleterious effects on the hippocampus.

